# Power analysis for human melatonin suppression experiments

**DOI:** 10.1101/2023.01.30.526303

**Authors:** Manuel Spitschan, Parisa Vidafar, Sean W. Cain, Andrew Phillips, Ben C. Lambert

## Abstract

In humans, the nocturnal secretion of melatonin by the pineal gland is suppressed by ocular exposure to light. In the laboratory, melatonin suppression is a convenient biomarker for this neural pathway. Recent work has found that individuals differ substantially in their melatonin-suppressive response to light, with the most sensitive individuals being up to 60 times more sensitive than the least sensitive individuals. Planning experiments with melatonin suppression as an outcome needs to incorporate these individual differences, particularly in common resource-limited scenarios where running within-subjects studies at multiple light levels is expensive and not feasible with respect to participant compliance. Here, we present a novel framework for virtual laboratory melatonin suppression experiments, incorporating a Bayesian statistical model. We provide a Shiny web app for power analyses that allows users to modify various experimental parameters (sample size, individual-level heterogeneity, statistical significance threshold, light levels), and simulate a systematic shift in sensitivity (e.g., due to a pharmacological or other intervention). Our framework helps experimenters to design compelling and robust studies, offering novel insights into the underlying biological variability relevant for practical applications.

## Introduction

Light exposure has a profound impact on human physiology and behaviour. In addition to enabling vision, light elicits a range of physiological and behavioural responses. These responses include the acute suppression of nocturnal melatonin production by light and shifting of the endogenous circadian rhythm (***Blume et al., 2019***; ***Vetter et al., 2021***). These effects, often summarised under the umbrella term ‘non-visual’ effects of light (***Schlangen and Price, 2021***), are mediated by a pathway connecting the eye to the hypothalamus. More specifically, the suprachiasmatic nuclei (SCN) receive retinofugal input, predominantly from a subset of retinal ganglion cells which are photosensitive in the absence of input from the cones and rods, the canonical photoreceptors underlying visual function. This photosensitivity is owed to the expression of the short-wavelength-sensitive photopigment melanopsin (***Provencio et al., 1998, 2000***; ***Do, 2019***; ***Spitschan, 2019***). There is converging and convincing evidence linking the spectral sensitivity of melanopsin to the *in vivo* sensitivity of circadian and neuroendocrine responses to light (***Brainard et al., 2001*** ; ***Thapan et al., 2001*** ; ***Nowozin et al., 2017*** ; ***Prayag et al., 2019*** ; ***Brown, 2020*** ; ***Gimenez et al., 2022***).

Most evidence for the melatonin-suppressive effects of light is generated in laboratory experiments, in which participants are exposed to carefully controlled illumination (for examples, see ***Cajochen et al. (2005***); ***Nagare et al. (2019***); ***Allen et al. (2018***); ***Spitschan et al. (2019***)), while their saliva, plasma, or urine is sampled for later melatonin assay (***Kennaway, 2019***, ***2020***). As these experiments can be resource-intensive in terms of participant burden, processing costs, and staff to run a specific multi-hour (and potentially multi-day) experimental protocols, an important consideration in designing experiments is statistical power.

Non-visual sensitivity to light has large individual differences (***Chellappa, 2021*** ; ***Spitschan and Santhi, 2021***). A recent study by ***Phillips et al. (2019***), in which participants were exposed to overhead illumination at different illuminance levels on different evenings while their melatonin production was measured, showed very large individual differences, with the most sensitive individual being up to 60 times more sensitive than the least sensitive individual in their data set. The sizeable individual differences require special attention in designing experiments to obtain compelling evidence, particularly when it is not feasible to run extensive within-subjects experiments sampling the same individuals multiple times under multiple illuminances (thereby minimising individual differences). Here, we present a novel framework for facilitating power analyses for human melatonin suppression experiments to optimize the choice of experimental illumance levels.

## Results

### Overview

We developed a Bayesian statistical model of a virtual laboratory melatonin suppression experiment (***Phillips et al., 2019***). The model can be thought of as representing the melatonin suppression versus photopic illuminance (“lux”) curves (henceforth termed “dose-response curves”) across the population from which the original study participants were sampled. The model is stochastic, meaning that each *virtual individual* drawn from it will likely have a different dose-response curve. Figure 1 shows four replicates of a *virtual experiment* comprising *n* = 41 participants in each case (the same sample size as the estimates in ***Phillips et al. (2019***)). There are two types of stochasticity inherent in the model: individual-level variation in dose-response curve parameters; and measurement error when recording an individual’s melatonin suppression level at a single discrete lux value. These two sources of uncertainty are evident in Figure 1 by the variation in dose-response curves between subjects and the variation in measurements (black points) around them.

**Figure 1.**
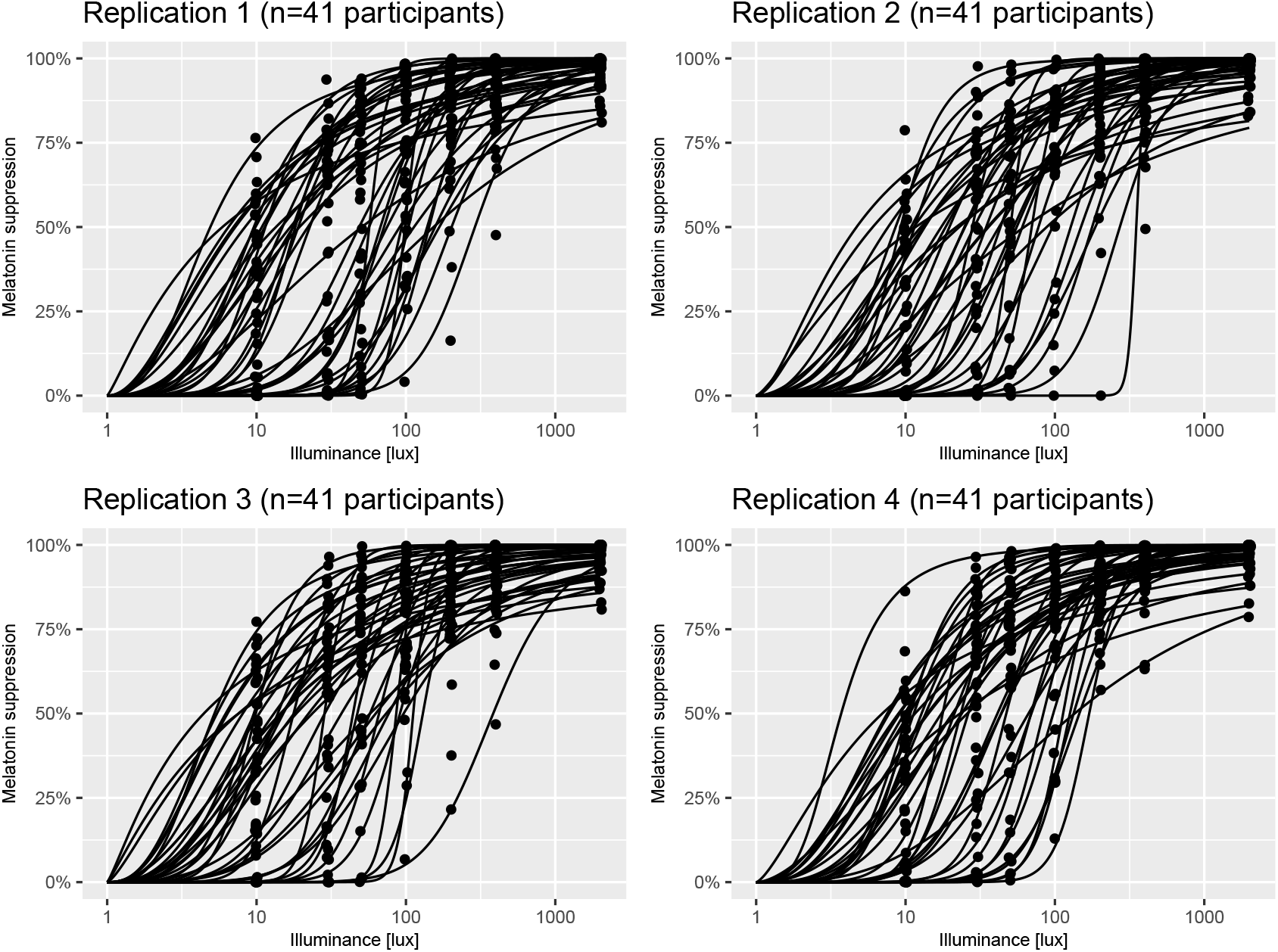
Example virtual experiments. Each panel shows individual dose-response curves and measurements (black points) of melatonin suppression at a series of lux values for *n* = 41 participants generated using the `virtual_experiment(n=41)` function from the `melluxdrc` R package. The assumptions underpinning each of these panels are described in *A model of virtual experiments*.

The assumptions underpinning our model of virtual experiments are detailed in *A model of virtual experiments*. The ability to generate such virtual experiments allows us to conduct a series of *in silico* experiments to estimate the statistical power for a series of different laboratory study designs. The various study types are outlined in §. As part of this work, we have created an opensource R package called *melluxdrc* (***Spitschan and Lambert, 2021d***) that wraps the functionality required to generate virtual experiments and to perform statistical power calculations. To facilitate power calculations for those designing experiments, we have also conducted a series of calculations and made those results freely available via an online R Shiny application called the *mellux-app* (***Spitschan and Lambert, 2021a***), which we describe further in §.

### A model of virtual experiments

Our model of virtual experiments comprises two elements: a) a population model of dose-response curves and b) a measurement error model that statistically represents the various noise factors that influence measurements of melatonin suppression at a particular lux level.

The population model of dose-response curves is based on parameter estimates presented for *n* = 41 participants in ***Phillips et al. (2019***). In ***Phillips et al. (2019***), dose-response data for *n* = 55 participants were obtained in a within-subjects protocol, where participants were exposed to a dim control (< 1 lux) and five other *experimental* light levels (10, 30, 50, 100, 200, 400, 2000 lux) for 5 hours in the evening. Using these data, melatonin suppression values, *s*(*x, i*), were obtained at each experimental light level, *x* for each participant *i*. The series of melatonin suppression values for each participant were then modelled using a logistic-type curve of the form:

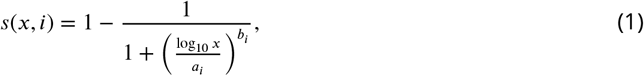

which has the property that melatonin suppression increases towards 100% as illuminance increases without bound; here, *b*_*i*_ > 0 controls the shape of each individual’s dose-response curve, and *a*_*i*_ = log_10_ ED_50_ for that individual. Here, *ED*_50_ is the dose corresponding to a melatonin suppression value of 50%.

The values used for our analysis were the (*a*_*i*_, *b*_*i*_) estimates for the *n* = 41 participants previously reported in which a reliable estimate of the dose-response curve was obtained (less than 1 log-unit 95% confidence interval for the estimate of *a*_*i*_): each set characterising the dose-response curve as in eq. (1). From here onwards, we refer to this set of estimates as the *raw dose-response estimates*. Using these values, we sought to create a statistical population model that represented these collection of dose-response curves which, crucially, could be sampled from to obtain a dose-response curve for a virtual individual.

To do so, we built a statistical distribution representing the raw estimates. To represent this bivariate distribution, we used kernel density estimation using the `bkde` function from the *KernSmooth* R package (***Wand, 2015***) to approximate the empirical distribution over *a*_*i*_ values. We then modelled the conditional distribution of log *b*_*i*_ conditional on *a*_*i*_ through a regression equation:

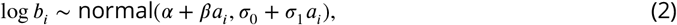

where the regression equation allows heteroscedasticity, represented by a standard deviation that increases linearly with *a*_*i*_ (since *σ*_0_ > 0 and *σ*_1_ > 0). This model was estimated using a Bayesian framework, meaning that priors were set on the parameters. The priors chosen were uninformative, allowing a wide range of possible relationships and are shown in Table 1. The model was fitted using Markov chain Monte Carlo (MCMC) through Stan’s NUTS algorithm (***Carpenter et al., 2017***), using 4 Markov chains with 4000 iterations per chain, with 2000 of each chain’s iterations discarded as warm-up; finally, the post-warm-up iterations were thinned by a factor of 2. MCMC convergence was diagnosed through 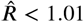 and having bulk- and tail-ESS values above 400 (***Vehtari et al., 2019***). Posterior predictive checks of model fit (see, for example, (***Lambert, 2018***)) indicated that the model was a reasonable fit to the data (Fig. 8). The Stan file and all materials needed to reproduce this analysis are available at (***Spitschan and Lambert, 2021b***).

**Table 1.** Priors. Shows the priors used on the parameters of the Bayesian models that were estimated.

| Model | Prior |
| --- | --- |
| eq. (1) | $\alpha \sim \text{normal}(0, 1)$ |
| eq. (1) | $\beta \sim \text{normal}(0, 1)$ |
| eq. (1) | $\sigma_0 \sim \text{Cauchy}^+(0, 1)$ |
| eq. (1) | $\sigma_1 \sim \text{Cauchy}^+(0, 1)$ |
| eq. (4) | $a \sim \text{Cauchy}(0, 1)$ |
| eq. (4) | $b \sim \text{Cauchy}(0, 1)$ |

To sample parameters characterising a dose-response curve for a virtual individual, we then used the approach described in Algorithm 1. Briefly, the first section of the algorithm draws *a* from the kernel density estimate of the empirical distribution. Then eq. (2) is used to draw a value of log *b* conditional on *a*. The final section of the algorithm calculates the *ED*_25_ and *ED*_75_ corresponding to the sampled *a* and *b* values. If either of these are more extreme than thresholds derived from the corresponding ED values from the estimates from (***Phillips et al., 2019***), those (*a, b*) values are rejected and the function is called again. This last step ensures that the dose-response curves obtained for virtual individuals are not far more extreme than those witnessed in the raw estimates. This approach is able to generate samples of (*a, b*) parameters that encompass the distribution of raw estimates (Fig. 9). Accordingly, the corresponding *ED*_25_ and *ED*_75_ values generated by this process were also a reasonable fit to those corresponding to the raw estimates (Fig. 9).

#### Algorithm 1 Virtual individual generation

Takes as input posterior draws of *α, β, σ*_0_, *σ*_1_ in eq. (2)

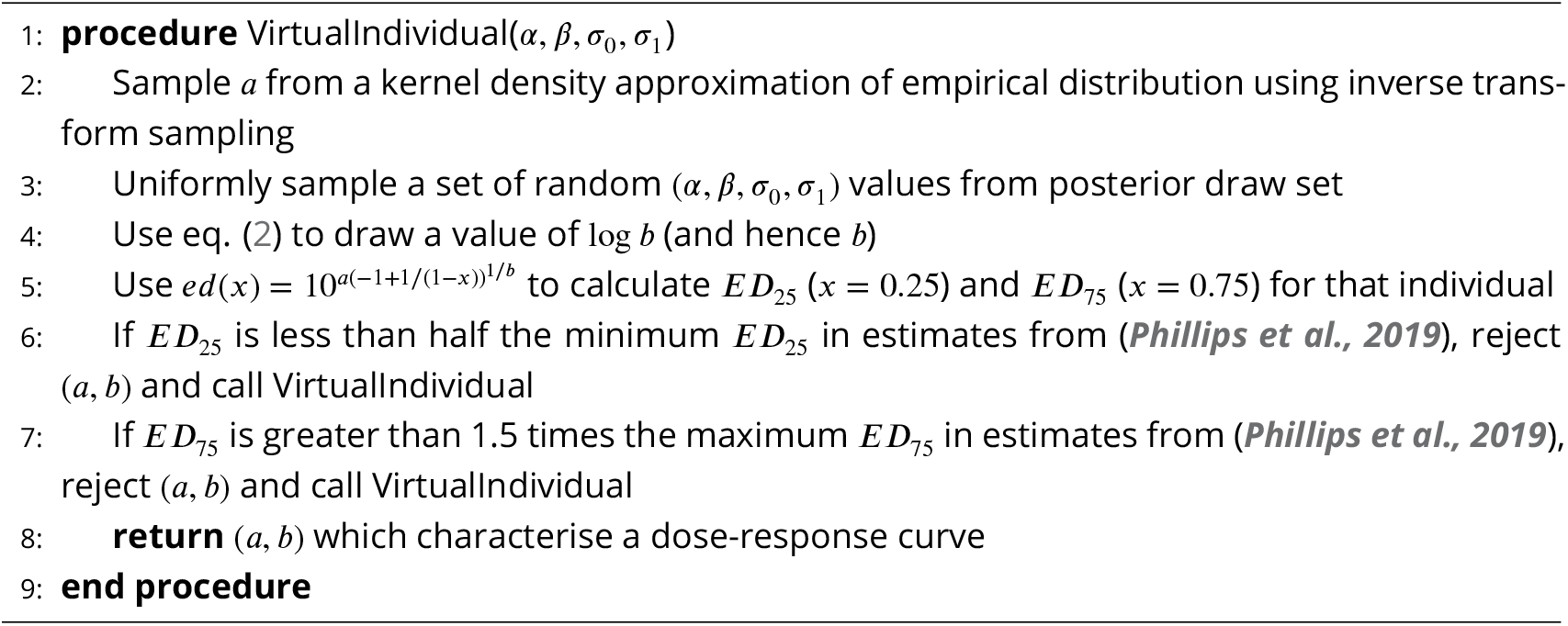

As part of our virtual population, we also developed an algorithm to allow users to simulate a reduction in individual heterogeneity in the population. To do so, we introduced a parameter, 0 ≤ *η* ≤ 1 that modulates the level of individual heterogeneity in the population: here, *η* = 0 indicates that virtual individuals sampled from the population all have the same dose-response curve near the population median; *η* = 1 indicates that the virtual individuals are drawn from the unrestricted population model (i.e., by the same process described in Algorithm 1). The process used to sample virtual individuals from the population is provided in Algorithm 2. Figure 2 shows how reducing the individual-level heterogeneity using this approach results in a tighter spread of dose-response curves.

**Figure 2.**
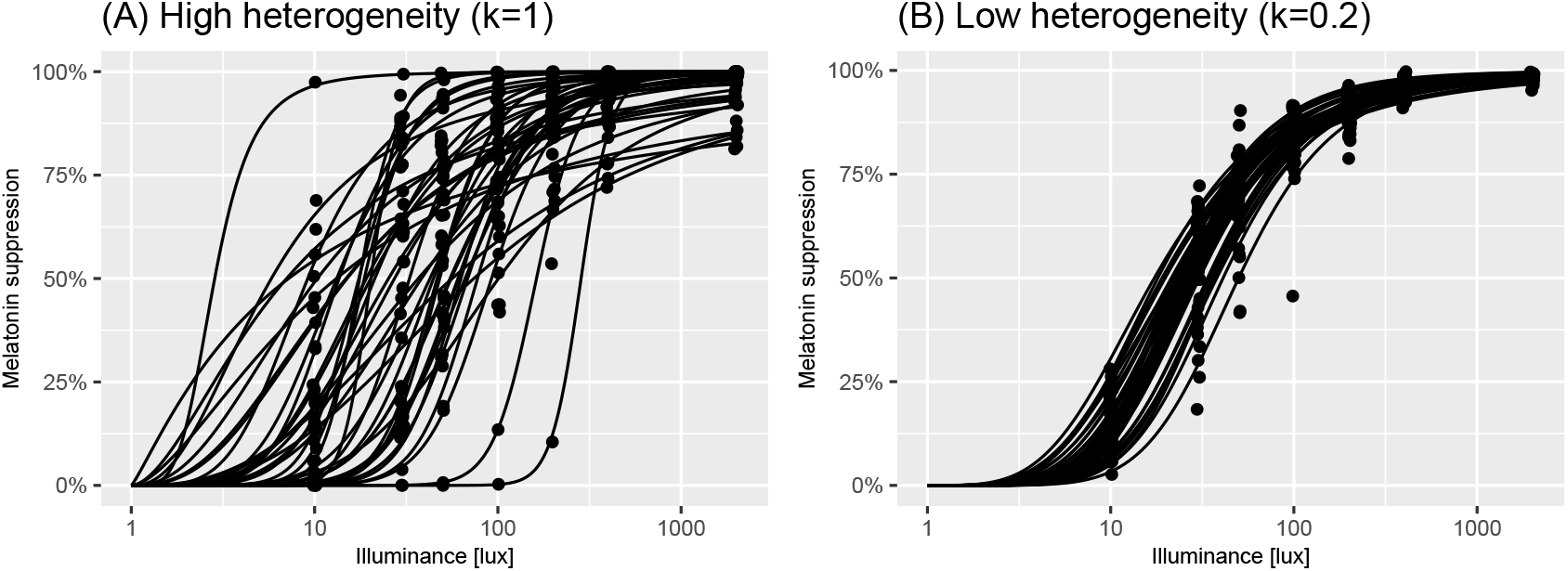
Effect of reduction in individual variance. Panel A shows individual dose-response curves and measurements (black points) of melatonin suppression at a series of lux values for *n* = 41 participants generated using the `virtual_experiment(n=41)` function from the `melluxdrc` R package. Panel B shows the same but assuming a reduction in individual variance given by *η* = 0.2 as given by the `virtual_experiment(n=41, individual_variation_level=0.2)`. The assumptions underpinning each of these panels are described in §.

#### Algorithm 2 Virtual individual generation: reduced individual variance

Takes as input posterior draws of *α, β, σ*_0_, *σ*_1_ in eq. (2) and *η*

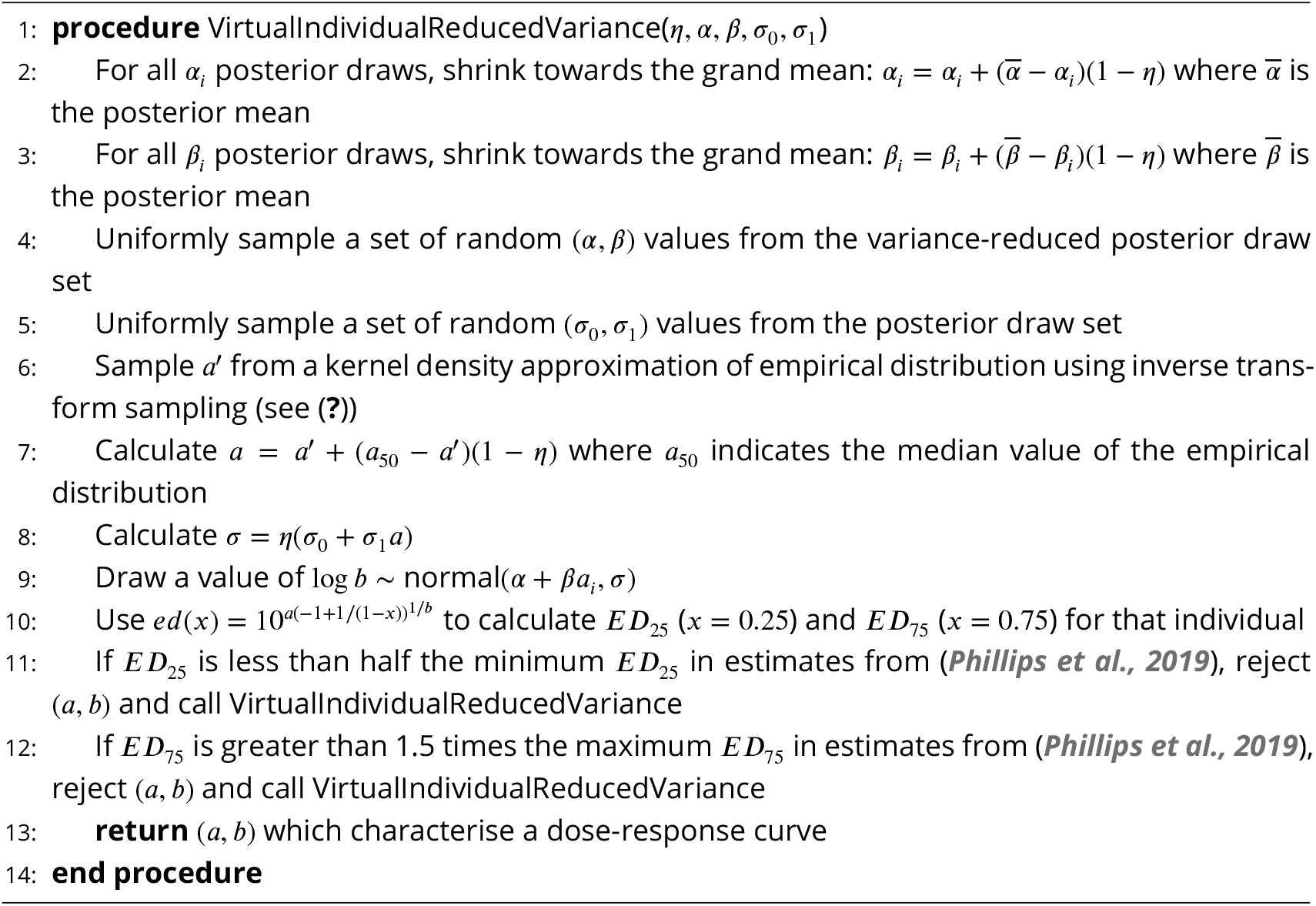

As is seen in Figure 2 in ***Phillips et al. (2019***), the measured melatonin suppression values at discrete lux values exhibited often considerable variation around the best fit lines. Using the Root-Mean-Square-Errors (RMSEs) for each fit in for each individual, we developed an error model representing measurement variability for each individual. To do so, we assumed that measurement noise was additive on the logit scale to ensure that measurements could not fall outside of the [0, 1] range. That is,

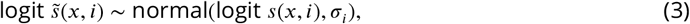

where logit(*z*) := log *z*/(1 − *z*), and 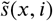 represents the measured melatonin suppression at lux *x* for individual *i*. Here, *σ*_*i*_ > 0 is an individual-specific value quantifying measurement noise.

We aimed to determine the value of *σ*_*i*_ that generated measurement noise resulting in a corresponding RMSE value close to that observed for individual *i*. To do so, we used an approximate estimation approach where, in each iteration, we simulated data using eqs. (1) and (3), and compared the simulated RMSE value to the truth. Specifically, we used the known (estimated) (*a*_*i*_, *b*_*i*_) values for individual *i* to simulate a dose-response curve given by eq. (1); we then generated measurements at the discrete lux values used in (***Phillips et al., 2019***) using eq. (3) with a particular *σ*_*i*_. For each such “experiment”, we calculated an RMSE value. For a given *σ*_*i*_ value, we repeated the experiment 100 times, and calculated an average RMSE value across all replicates. The difference between this average simulated value and the true RMSE value was then used as the target for the one-dimensional root-finding algorithm `uniroot` available in R: the output of this algorithm was, hence, a value of *σ*_*i*_ for each individual.

In order to generate measurements for virtual individuals, we needed an approach to generate potential *σ*_*i*_ values for unseen individuals. To do so, we assumed that the individual *σ*_*i*_ values were drawn from the following population process:

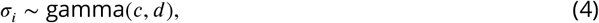

where *c* > 0 and *d* > 0. We estimated the parameters in eq. (4) using the values of *σ*_*i*_ estimated by the root-finding algorithm. The model was estimated in a Bayesian paradigm, requiring that we set priors on the parameters. Here, we chose uninformative priors on the parameters, which are specified in Table 1. The model was fitted using Markov chain Monte Carlo (MCMC) through Stan’s NUTS algorithm (***Carpenter et al., 2017***) using 4 Markov chains with 2000 iterations per chain, with 1000 of each chain’s iterations discarded as warm-up. The same criteria were used to diagnose model convergence as for eq. (2). Posterior predictive checks of model fit indicated that the model was a reasonable fit to the data (Fig. 8).

With both elements of the virtual experiment model in place – the population model of dose-response experiments and the measurement error model – we sought to assess how well the resultant model represented the data collected in (***Phillips et al., 2019***). This was tested using the minimum and maximum melatonin suppression at each measured lux value (the “extrema”); and the percentage of observations which were extreme at each lux: either below 5% lux or above 95% melatonin suppression (the lower and upper “saturation values”). We generated 200 replicate virtual experiments, which each had the same number of individuals (*n* = 41) as the estimates we were provided with. For each experiment and lux value, we calculated simulated extrema and saturation values to compare with the real measurements. In Figure 9, we compare the modelled extrema (black lines) with the real values (orange points and line): in the left-hand plot, we compare the minima; in the right plot, we compare the maxima. This indicates that, for moderate-high lux values, the modelled and real values were in good accordance. At the lower lux values, the modelled values tended to be more extreme (either closer to 0% for the minima; or closer to 100% for the maxima) than the real data. In Figure 9, we performed the same comparison but for the levels of simulated and actual saturations. In this case, the upper saturation values between the modelled and real data were reasonable, apart from at the highest lux value measured. The lower saturations exhibited the same pattern as for the extrema, in that the simulated data were more extreme than the actual.

The discrepancies between the modelled and actual data could be due to a) assumptions around our model for the population of dose-response curves, b) assumptions around our measurement process model and/or c) issues with the original logistic models used to model dose-response curves in (***Phillips et al., 2019***). Our checks for our model of dose-response curves (Fig. 9) indicate no issue with representing the original estimates. It is possible that the logit-normal noise process we assume in eq. (3) results in greater variation than seen in reality. Visual inspection of Figure 2 in (***Phillips et al., 2019***), however, shows that, for a number of individuals, the logistic dose-response curve is downward-biased for low illuminance levels, potentially indicating that a two-parameter logistic is inappropriate in this extreme. Without the raw experimental data, however, it is not possible to determine the exact cause of the discrepancy between modelled and real life data. Despite these differences, the overall correspondence between the model and data is good, and, as such, it provides a reasonable basis to determine statistical power for a variety of different experimental settings.

### Power calculations

Having developed a virtual experiment model (see *A model of virtual experiments*), we sought to use it to help inform experimental design. In particular, we considered two classes of experiment: in *lux comparison experiments*, the aim is to quantify whether population melatonin suppression differs across two measured lux values; in *intervention experiments*, the aim is to estimate the effect size for an intervention that systematically changes the dose-response curves (i.e., changes light sensitivity). Within each experimental class, we considered two possible experimental designs: within-subject designs, where the same participants are measured twice; or between-subject designs, where two separate groups of participants are measured once each. In all cases, we used our model of virtual experiments to generate data that mimics the types of laboratory experiments described here, which was then used to determine statistical power.

In lux comparison experiments, individual melatonin suppressions are measured at two lux values: *x*_1_ and *x*_2_. In within-subject experiments, each individual is measured twice resulting in *N* paired observations: 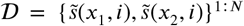. In between-subject experiments, there are 2*N* individuals, with *N* measured at *x*_1_ and another *N* at *x*_2_, resulting in a dataset: 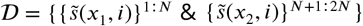. In these experiments, the aim is to determine that there is a significant difference in the suppression response between measurements taken at *x*_1_ and *x*_2_. To do so, we use a t-test which is either paired (for within-subject designs) or independent (for between-subject designs). We considered all possible pairs of different lux values from the set {10, 30, 50, …, 1970, 1990}. For each pair, we also considered a range of sample sizes from *N* = 10, 20, …, 90, 100 and a range of possible individual heterogeneity values: *η* = 0, 0.2, 0.4, 0.6, 0.8, 1.0. For each combination of these parameters, we calculated the power to detect significant differences of the correct sign using test sizes of 1%, 5% and 10%.

We show visualisations of the outcomes for these analyses in Figures 4 and 5. Figure 4 shows the statistical power as a function of participant level hetereogeneity for between- and within-subjects designs. The same principle of power but for different illuminances levels is shown in 5. The results are clear: within-subjects designs beat between-subjects designs.

**Figure 3.**
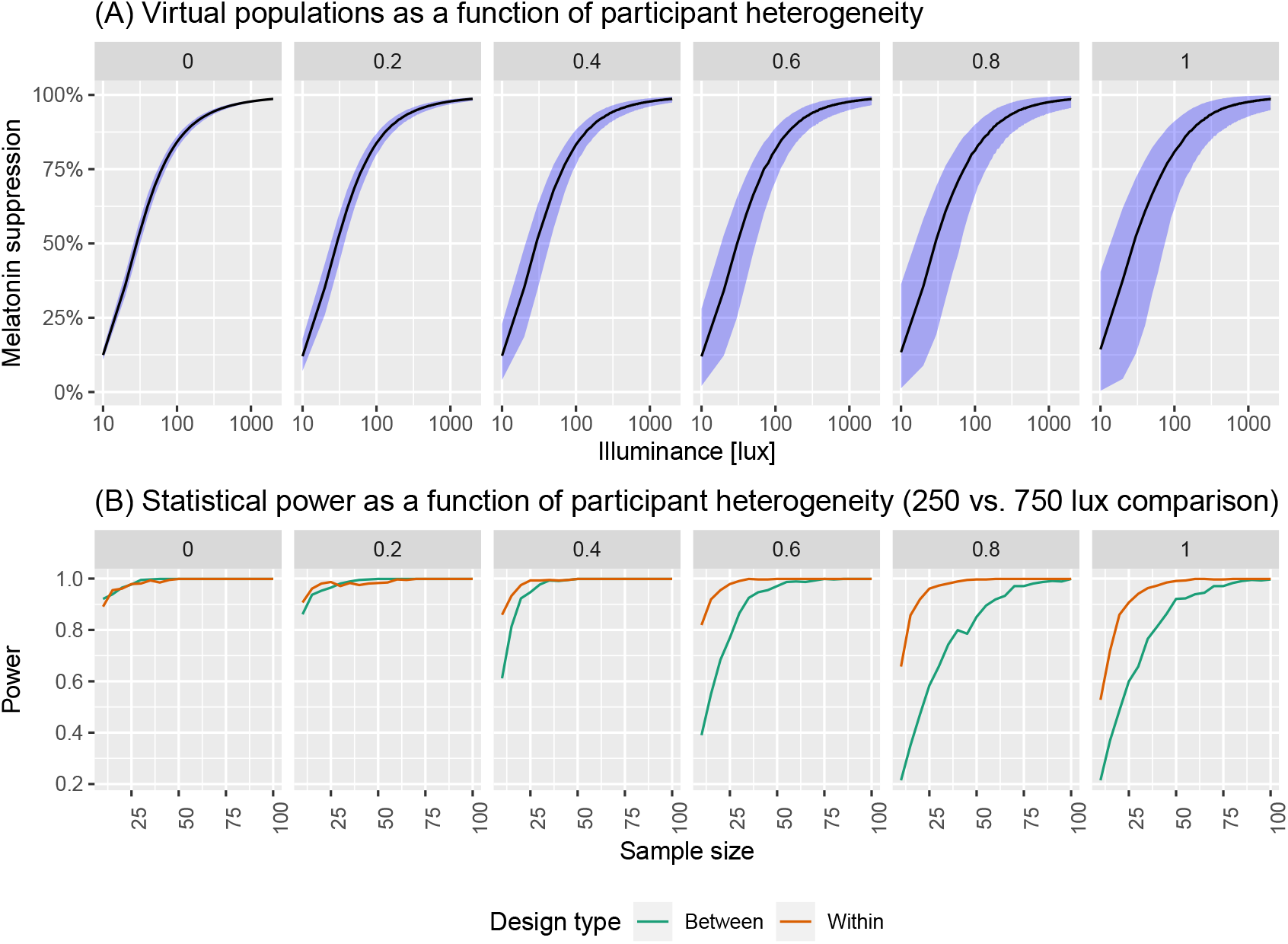
(A) Virtual populations as a function of participant heterogeneity. Here, we are simulating the group dose-response curves assuming different levels of participant heterogeneity. (B) Statistical power as a function of participant heterogeneity.

**Figure 4.**
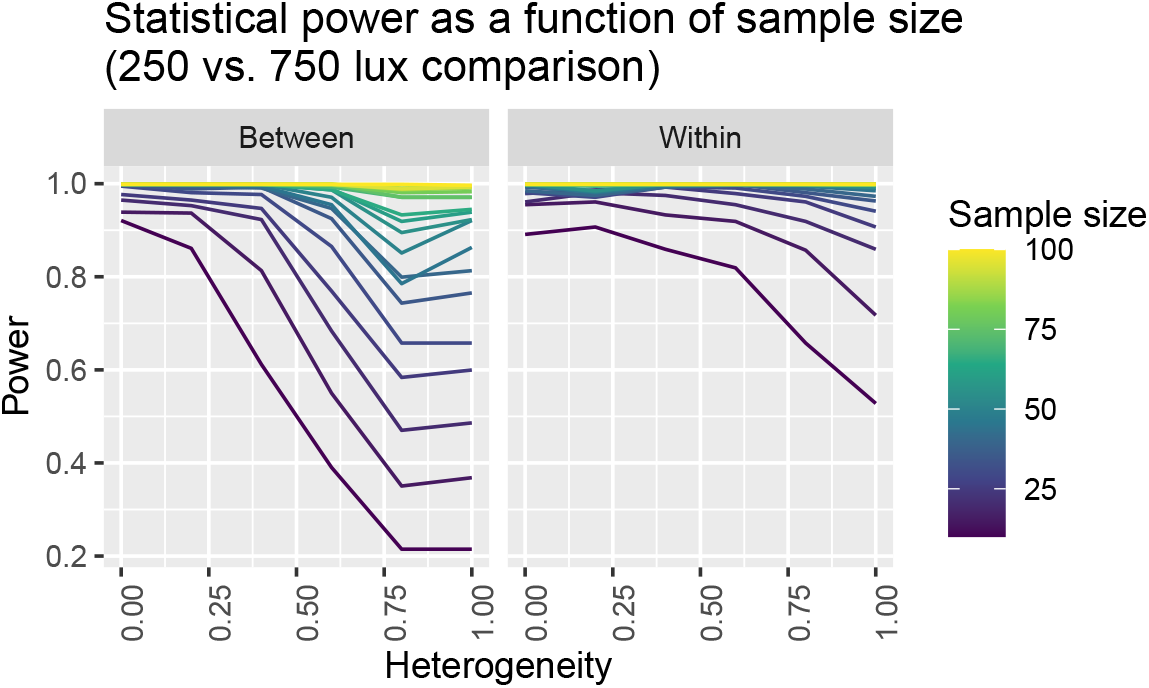
Statistical power for detecting a difference in melatonin suppression assuming different samples sizes and between-subjects (left panel) and within-subjects (right panel) designs.

**Figure 5.**
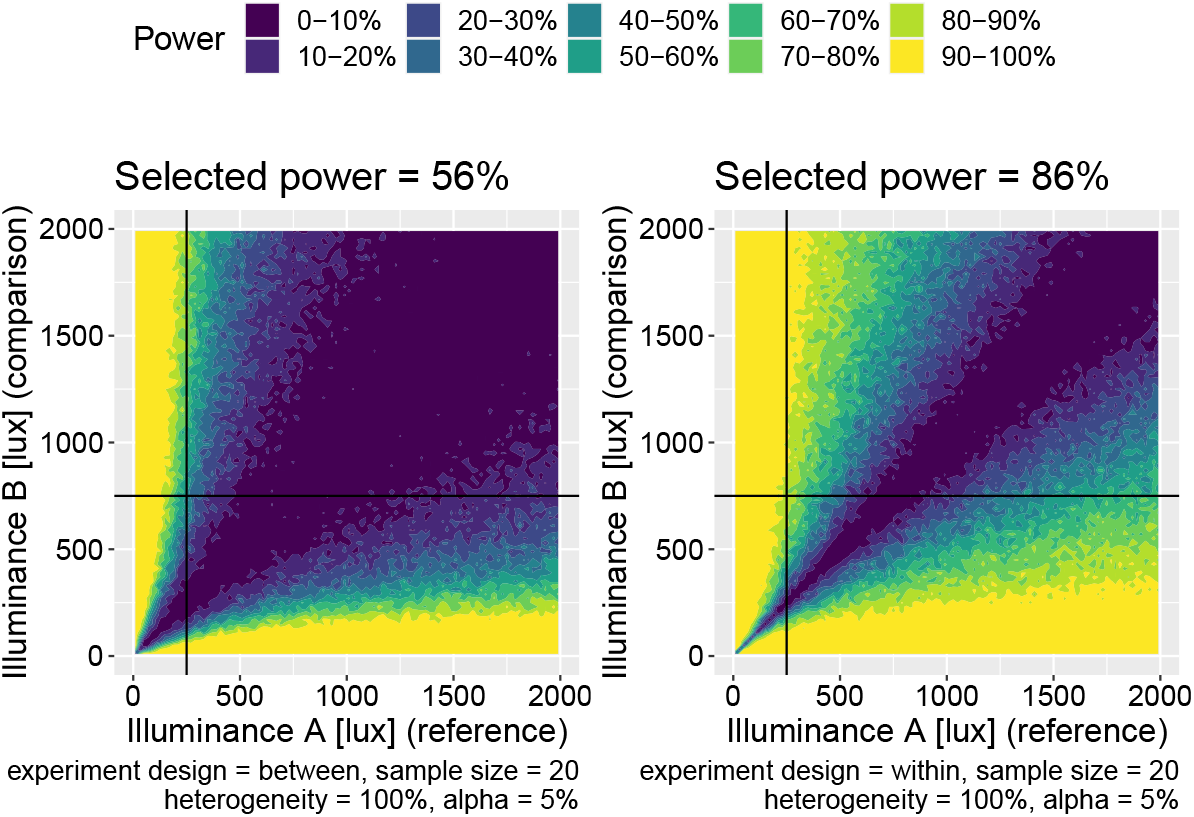
Power maps for two-lux comparison experiments for between-subjects (left panel) and within-subjects experiments (right panel). The choice of comparison light levels here is 250 and 750 lux.

**Figure 6.**
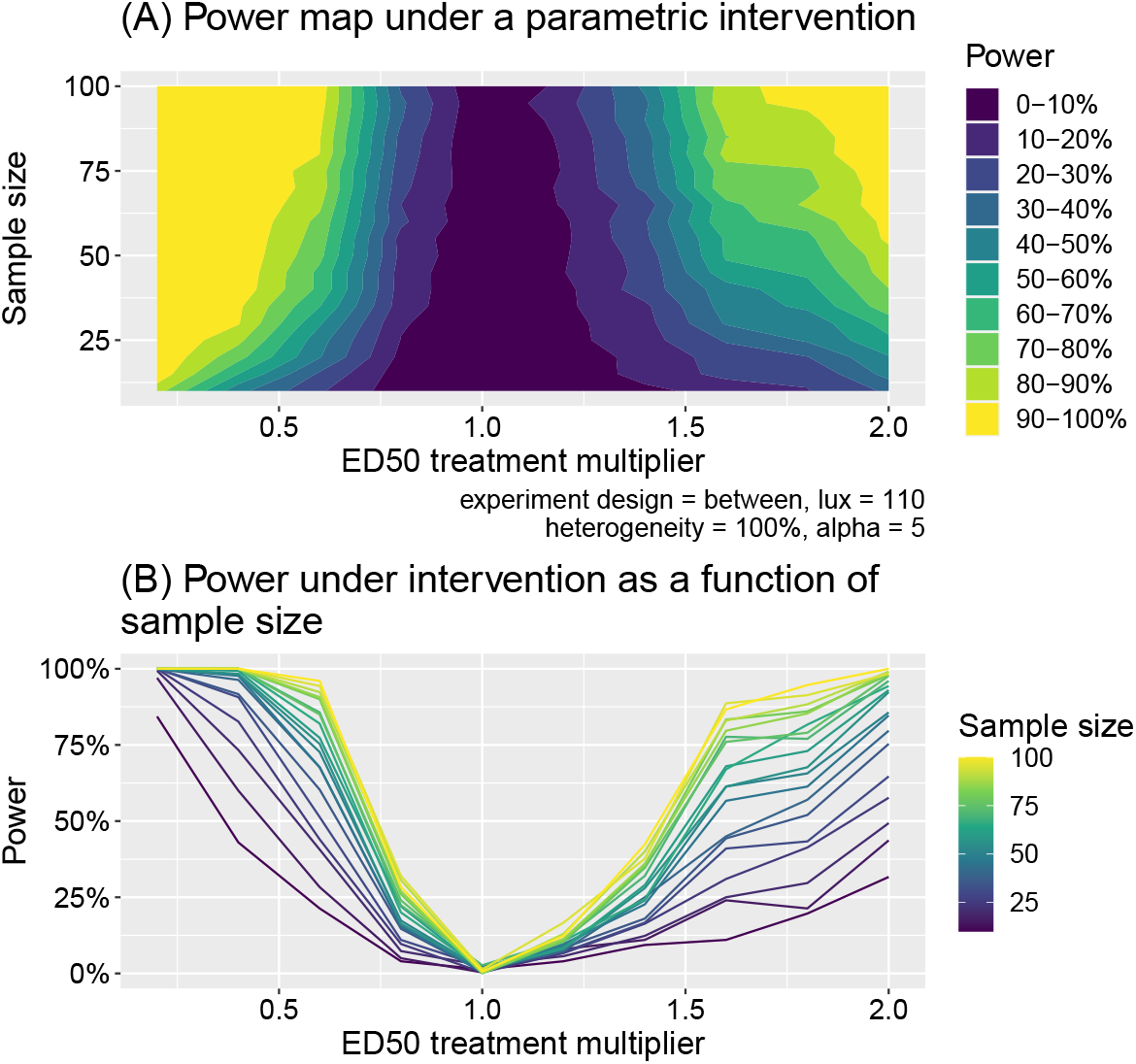
Panel A shows the power as a function of the intervention effect (expressed as a multiplier of the ED50 value) and sample size. Panel B shows a slice of the same data as a line plot for different discrete samples sizes.

**Figure 7.**
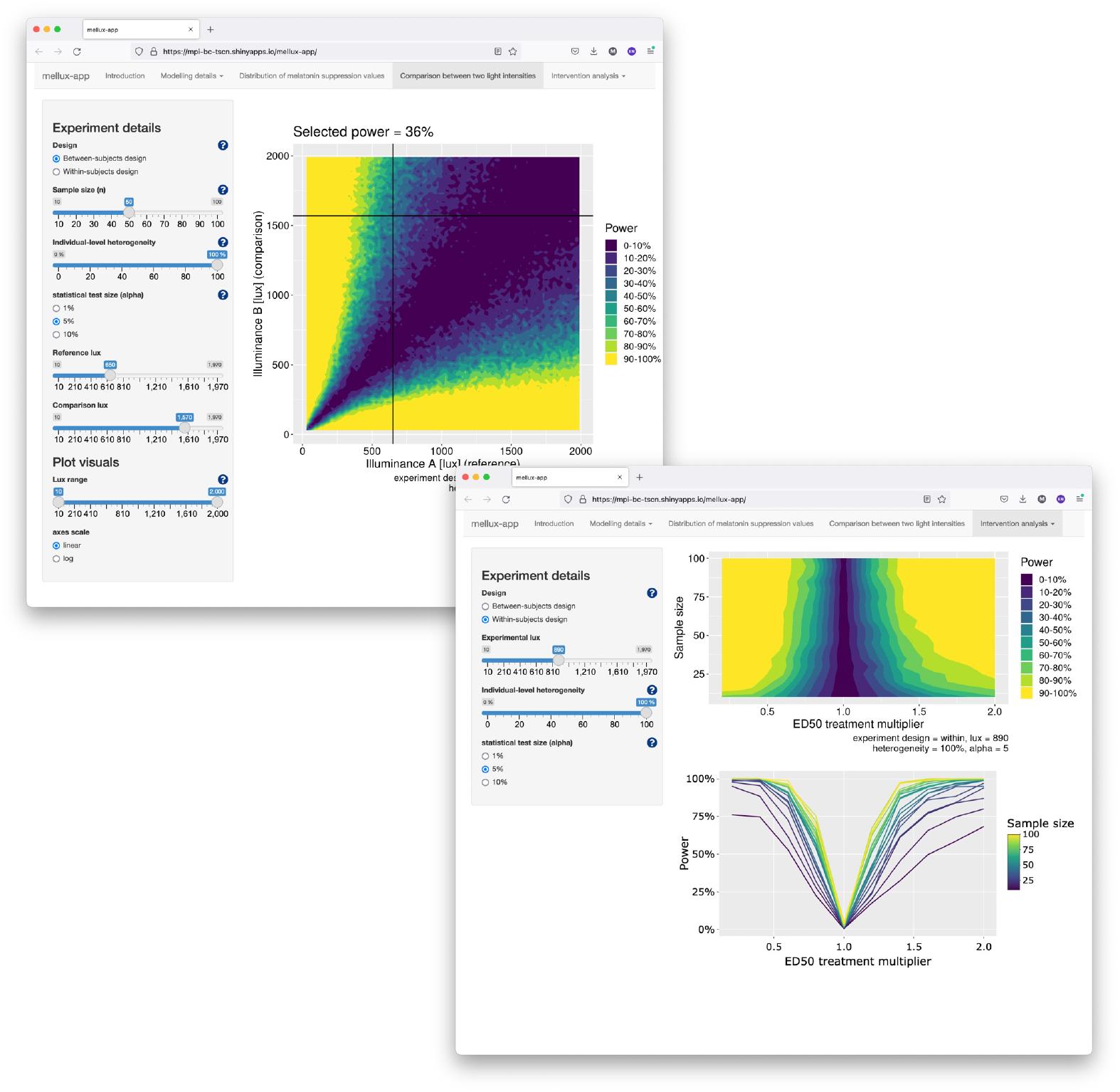
Screenshots of the Shiny app for exploring sample size calculations interactively.

In intervention experiments, we assume interventions that shift the ED_50_ away from the baseline level. In within-subject study designs, individuals are assumed to have their melatonin suppressions measured at a particular lux *x* once before the intervention occurs and once afterwards, resulting in a dataset: 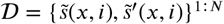 where unprimed variables indicate natural levels and primed indicate intervention levels. For the between-subject design, two different sets of individuals were measured: a “baseline” group and an “intervention” group whose ED_50_ values are shifted. This resulted in a dataset 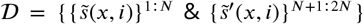. As for the lux comparison experiments, we used t-tests to compare the natural and intervention melatonin suppressions: again using paired-tests for the within-subject design and independent tests for the between-subject design. We considered all lux values from the set *x* = {10, 30, 50, …, 1970, 1990} across the same set of sample sizes and individual heterogeneities as for the comparison experiments. Additionally, we considered natural ED_50_ multipliers taking values *χ* = {0.2, 0.4, …, 1.8, 2.0}, where, for example, *χ* = 0.2 means the natural ED_50_ of individuals are reduced by a factor of 5. For each combination of these parameters, we calculated the power to detect significant differences of the correct sign using test sizes of 1%, 5% and 10%.

We summarise the outcomes of these analyses in 6. We find clear differences in power assuming a specific intervention effect. These analyses may serve to inform sample size calculations for planning interventions affecting the impact of light on melatonin suppression.

## Conclusion

Here, we have developed a model of virtual melatonin suppression experiments under different light levels informed by, and fitting to, actual experimental data. We expanded our model virtual experiments by allowing the modulation of individual-level variability and implementing a virtual intervention leading to a left- or rightward shift of the dose-response curve. We provide a web applet for exploring the virtual model and using it in practical applications.

## Methods

### Reproducibility of results

The version of `melluxdrc` used to generate the results in this paper is v1.0.0.

### Shiny app

We provide an app developed in the Shiny framework which allows users to generate power analyses. The app is deployed at https://mpi-bc-tscn.shinyapps.io/mellux-app/. Screenshots are shown in 7.

## Code and data availability

All code and data underlying this work is available under the MIT License. We make available the R package `melluxdrc` for generating virtual experiments (***Spitschan and Lambert, 2021d***), code to run simulations on a cluster (***Spitschan and Lambert, 2021c***), and code to fit dose response curves (***Spitschan and Lambert, 2021b***). The code underlying the shiny app, deployed at https://mpi-bc-tscn.shinyapps.io/mellux-app/ is available as well (***Spitschan and Lambert, 2021a***).

## Supporting Information

**Figure 8.**
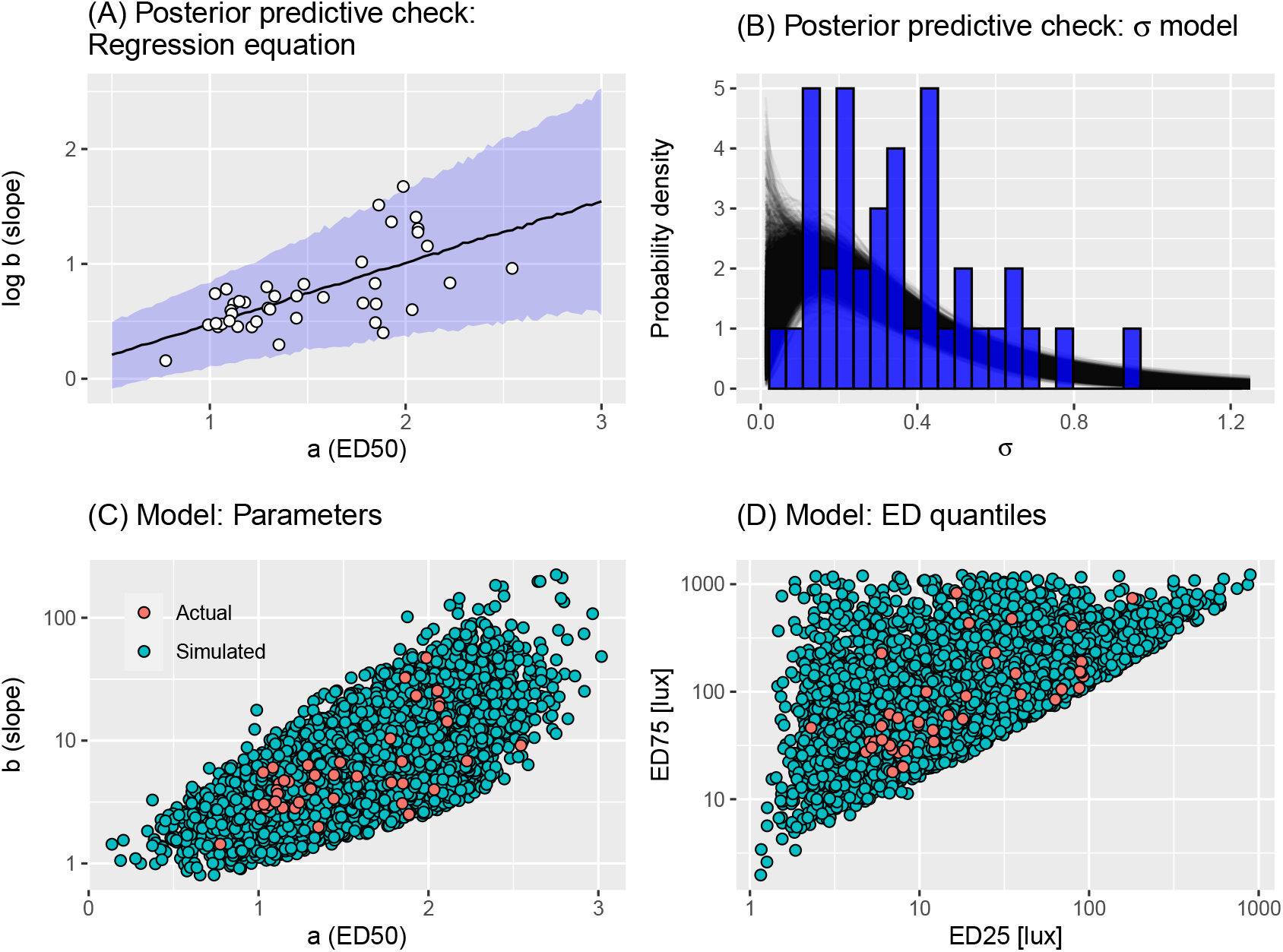
**(A) Posterior predictive check: regression equation.** Plot shows a graphical check of the model fit of eq. (2). The individual points show the estimates of (*a*_*i*_, *b*_*i*_) parameters provided by the authors of ***Phillips et al. (2019***). The uncertainty ribbon indicates the 2.5%-97.5% posterior predictive quantiles; the black line indicates the 50% posterior predictive quantile. **(B) Posterior predictive check:** *σ*_*i*_ **model**. Plot shows a graphical check of the model fit of eq. (4). Each black line represents a gamma density function corresponding to particular posterior samples of the parameters. The blue bars indicate the values of *σ*_*i*_ estimated by the root-finding algorithm. **(C) Assessing virtual individual generation: dose-response parameters**. Each orange point represents a draw of (*a, b*) parameters (in eq. (1)) obtained via Algorithm 1: here, we show 25,000 such estimates; each green point represents raw estimates from (***Phillips et al., 2019***). **(D) Assessing virtual individual generation:** *ED*_*x*_ **quantiles**. Each orange point represents the (*ED*_25_, *ED*_75_) values correspond to a draw of (*a, b*) parameters (in eq. (1)) obtained via Algorithm 1: here, we show 25,000 such estimates; each green point represents the *ED*_*x*_ quantiles corresponding to the raw estimates from (***Phillips et al., 2019***).

**Figure 9.**
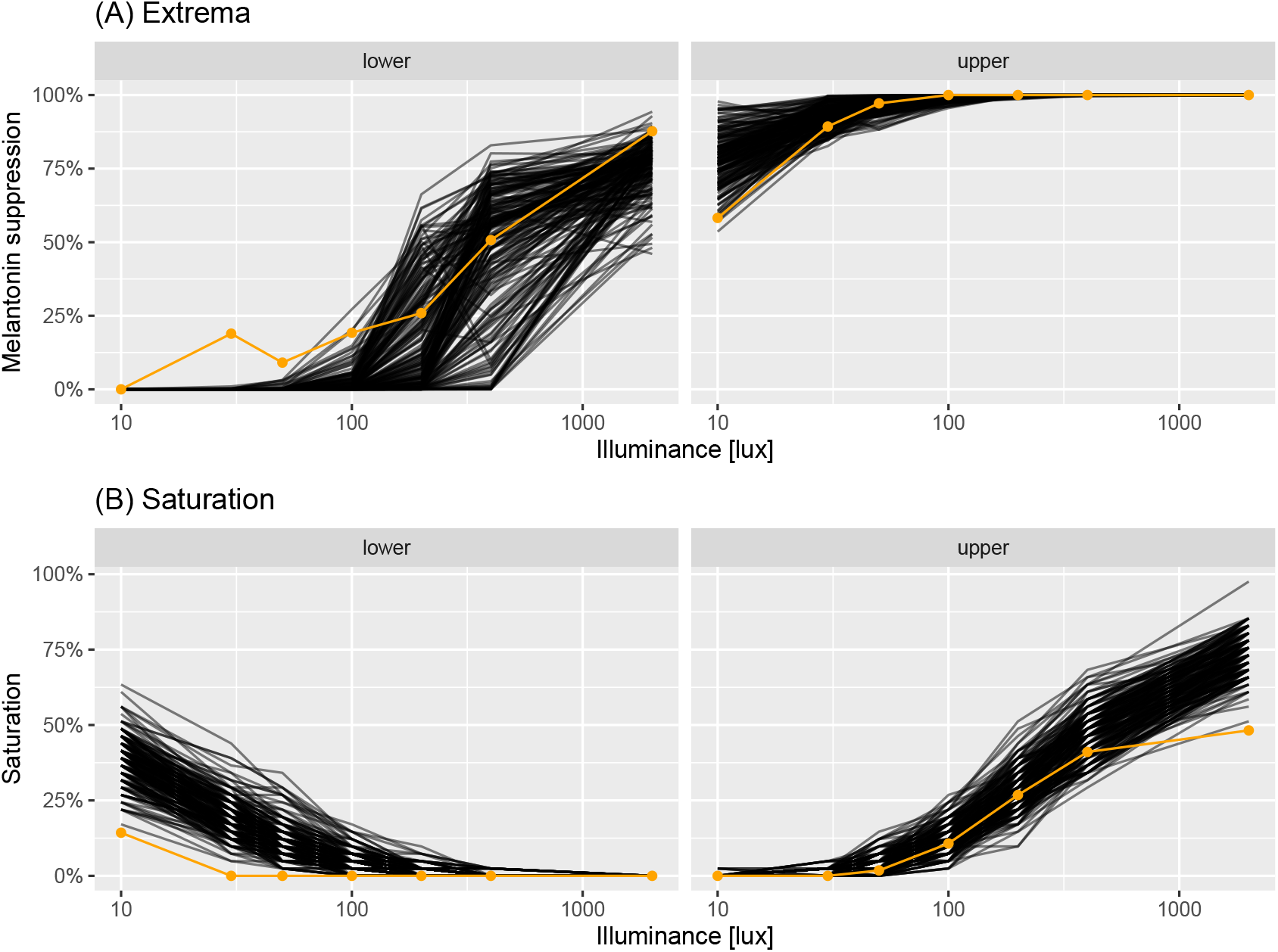
Virtual experiment check: saturations. Panels correspond to lower (percentage of observations < 5%) and upper (percentage of observations > 95%) saturations. Each black line corresponds to saturations generated from a single virtual experiment of sample size *n* = 41. Each orange point corresponds to the real saturation.

